# Excess fermentation and lactic acidosis as detrimental functions of the gut microbes in treatment-naive TB patients

**DOI:** 10.1101/2023.12.05.570051

**Authors:** Milyausha Yunusbaeva, Liliya Borodina, Darya Terentyeva, Anna Bogdanova, Aigul Zakirova, Shamil Bulatov, Radick Altinbaev, Fanil Bilalov, Bayazit Yunusbayev

## Abstract

Gut microbiota link to host immunity motivated numerous studies of the gut microbiome in tuberculosis (TB) patients. However, these studies did not explore the metabolic capacity of the gut community, which is a key axis of impact on the host’s immunity. We used deep sequencing of fecal samples from 23 treatment-naive TB patients and 48 healthy donors to reconstruct the metabolic capacity and strain/species-level content of the gut microbiome. We show that the strong taxonomic divergence of the gut community in TB patients is explained by the systematic depletion of the commensal flora of the large intestine, *Bacteroidetes,* and an increase in *Actinobacteria, Firmicutes*, and *Proteobacteria* such as *Streptococcaceae, Erysipelotrichaceae, Lachnospiraceae,* and *Enterobacteriaceae.* The cumulative expansion of diverse disease-associated pathobionts in patients reached 1/4 of the total gut microbiota, suggesting a heavy toll on host immunity along with MTB infection. Reconstruction of metabolic pathways showed that the microbial community in patients shifted toward rapid growth using glycolysis and excess fermentation to produce acetate and lactate. Higher glucose availability in the intestine likely drives fermentation to lactate and growth, causing acidosis and endotoxemia. Excessive fermentation and lactic acidosis are likely characteristics of TB patients’ disturbed gut microbiomes. Since lactic acidosis strongly suppresses the normal gut flora, directly interferes with macrophage function, and is linked to mortality in TB patients, our findings highlight gut lactate acidosis as a novel research focus and a potential host-directed treatment target that can augment traditional TB treatment.

## Introduction

Tuberculosis (TB), caused by *Mycobacterium tuberculosis*, is one of the leading infectious disease killers worldwide. In 2021, according to the World Health Organization (WHO), 10.6 million people were diagnosed with tuberculosis, an increase of 4.5% from 2020 (Global Tuberculosis Report 2021, 2021) Host immunity plays a central role in combating *M. tuberculosis* (MBT). Despite progress in elucidating the key elements of the innate (Liu et al., 2017) and adaptive immune responses to TB infection (Dheda et al., 2010; Scriba et al., 2017) host factors governing susceptibility to infection are poorly understood. It is unclear why some individuals in TB-endemic countries never become infected with MBT or why most latently infected individuals never progress to active TB disease. Among factors contributing to host susceptibility, the intestinal microbiota is gaining attention due to its prominent role in modulating host immunity. There is growing evidence that the gut microbiota plays a key role in developing the host immune system (Liu et al., 2017; Hooper et al., 2012) In connection to tuberculosis, several recent studies have attempted to assess the potential role that the gut microbiota can play in the host’s susceptibility to TB, both in humans and in model organisms.(Majlessi et al., 2017; Cardona et al., 2015). Specifically, experiments using model organisms suggest that gut microbiota can modulate host immunity and that microbiome changes can impact TB’s outcome and prognosis (Negi et al., 2020; Round & Mazmanian, 2010; Winglee et al., 2014). Despite these insights from model organisms, our understanding of the gut community in TB patients is limited to descriptive data on taxonomic alterations. Crucially, the functional impact of the microbiota on host immunity, mediated through microbial metabolic output, is lacking. Thus, based on 16S rRNA sequencing, earlier studies reported gut dysbiosis and discussed the possible impact of the detected taxonomic changes on the host (Hu et al., 2019; Shi et al., 2021; S. Wang et al., 2022; Yu et al., 2023). Such taxonomic descriptions are not enough to understand microbial function. There is increasing appreciation that gut microbes impact the human host more frequently via metabolic products (Rooks & Garrett, 2016). Therefore, insight into the metabolic capacity of the gut community is important to evaluate the potential effect of the microbiome on host immunity.

In this study, we used deep shotgun sequencing of fecal microbiomes in 23 treatment-naive TB patients and 48 healthy donors to dissect not only “who is present” in the gut community at the species/strain level but also infer “what they can do” by constructing metabolic functions from microbial genes. We showed that strong alterations of the gut communities in TB patients are accompanied by strong shifts toward metabolism indicative of microbial cell division and growth. Increased bacterial turnover in patients is accompanied by an increased capacity for glycolysis and fermentation of glucose to produce acetate and lactate. Of note, lactic acid production was the top change in TB patients’ guts, and lactic acidosis is known to increase lung pathology and mortality. Recent findings explain this detrimental effect by direct interference of lactic acid with macrophage activity against MTB and increased tissue destruction that worsens pathology. Thus, our findings came just in time to highlight the research focus on gut-derived lactic acidosis in tuberculosis and, perhaps, borrow treatment strategies from the field of animal husbandry, where this issue is well studied.

## Methods

### Study design and patient information

Our work is a case-control study with 23 treatment-naive TB patients and 48 healthy controls (HCs). TB patients were recruited at the Republican Clinical TB Dispensary (Ufa), Russia, during 2019–2021. All patients had newly diagnosed active pulmonary TB as confirmed by chest radiography and a positive sputum smear or were positive for *M. tuberculosis* based on the GeneXpert MTB/RIF test without evidence of rifampin resistance. Exclusion criteria included serious comorbidities, hemoptysis, hypoxia, extrapulmonary tuberculosis, history of tuberculosis treatment, use of antituberculous drugs within the past 30 days, pregnancy or breastfeeding, and HIV infection.

Participants for the control group (n = 48) were recruited from staff working at the Institute of Biochemistry and Genetics of the Russian Academy of Science (Ufa, Russia), considering the matched age and sex of the enrolled TB cases. These control subjects had no unexplained symptoms or other medical conditions, especially infectional, autoimmunity, or gastrointestinal diseases. All subjects had no change in chest radiography.

We collected freshly frozen samples from 48 healthy adult donors and 23 treatment-naive TB patients. Fecal samples from TB patients were taken before treatment started. All participants signed an informed consent form prior to entering the study.

### Fecal sample collection and DNA extraction

Fresh stool samples from the participants were collected using sterile containment and stored at 80 °C immediately for further analysis. Metagenomic DNA was extracted from 200 mg of feces using the QIAamp DNA Stool Mini Kit (QIAgen, Netherlands).

### Metagenomic Sequencing

Sequencing libraries were generated using the TruSeq DNA PCR-Free Sample Preparation Kit (Illumina, USA), following the manufacturer’s recommendations. The sequencing libraries were then sent for sequencing using the Illumina HiSeq2500 instrument, and at least 15 million 150 bp paired-end reads were generated for each donor.

### Raw sequence QC and preprocessing

We examined sequence quality before and after quality trimming and decontamination using FastQC v0.11.9 (Wingett & Andrews, 2018). Trimmomatic version 0.33 (Bolger et al., 2014) was used to remove low-quality bases (Q20), and fastp v0.23.2 was used to clip Illumina adapters (Chen et al., 2018). We additionally removed tandem repeats using Tandem Repeats Finder Version 4.09 (Benson, 1999). Finally, the KneadData v0.10.0 pipeline of the bioBakery toolset was used to decontaminate reads originating from the human genome, transcriptome, and microbial RNA (McIver et al., 2018). Specifically, we used the human reference genome (build hg37), the human reference transcriptome (build hg38), and the SILVA ribosomal RNA reference databases.

### Bacterial community diversity and principal component analysis

Shannon alpha and beta diversity measures were estimated using the microbiome R package (Leo Lahti, 2017). Differences in Shannon alpha diversity between patients and controls were tested using the two-sample Kolmogorov-Smirnov test implemented in the ks.test() function of the stats R package (R Core Team, 2022). To visualize gut community differences between donors using major axes of microbial variation, we applied principal component analysis (PCA) on the species abundance matrix. To adapt sparse data on relative abundance for principal component analysis, we imputed zeros in the relative abundance table using the cmultRepl() function in the zCompositions R package version 1.4.0-1 (Palarea-Albaladejo & Martín-Fernández, 2015). Next, to take compositionality into account, the relative abundance table was transformed using the center log ratio (CLR) prior to PC analysis. PC analysis was carried out using the prcomp() function in the stats R package (R Core Team, 2022).

### Inference of taxonomic content of the gut microbiome and between-group comparisons

Prior to downstream analyses, all bacterial taxa observed only in a single donor and at a relative abundance less than 0.0001 were discarded. The choice of an abundance threshold ensured that we kept rare bacterial species that can impact human disease, as demonstrated in recent work (Chriswell et al., 2022). Metaphlan v4.0 tool was used to infer both known bacterial species and strains as well as uncharacterized bacterial species. Uncharacterized bacterial species in the metaphlan v4.0 tool are defined in terms of species-level genome bins (SGBs) using so-called metagenomically assembled genomes (MAGs) (Blanco-Míguez et al., 2023). To facilitate comparison with published data, we analyzed relative abundances of bacterial taxa in sampled groups (TB patients and controls) at different taxonomic levels by aggregating strains and species into genus, family, order, class, and phylum. We used linear discriminatory analysis with effect size estimation (LEfSe) implemented in the microbial R package to identify bacterial taxa that characterize differences between patients and controls. We also computed the average proportion of each taxon between patients and controls at different levels (e.g., phylum and class). A statistical difference in proportions between groups was tested using the ANCOM statistical framework (Mandal et al., 2015) using the FDR (false discovery rate) approach to adjust p-values. When we tested average proportions using ANCOM, we were interested in comparing dominant taxa between groups. Therefore, we required bacterial taxa to be present in 25% of donors at an abundance of 5% or more.

### Inference of the microbial metabolic capacity of the gut microbiome and between-group comparisons

The metabolic capacity of the microbiome, often referred to as the metabolic profile (potential), was inferred using the HUMAnN 3.6 framework using default settings. Briefly, HUMAnN 3.6 compares the sequences from donor metagenomes to known microbial genomes and genes to infer known metabolic reactions and aggregate them into known microbial pathways. Specifically, for bioinformatic queries, HUMAnN 3.6 uses the ChocoPhlAn 3 database of 99.2 thousand annotated reference genomes from 16.8 thousand microbial species in the UniProt database (January 2019) (UniProt Consortium, 2019) and the corresponding functionally annotated 87.3 million UniRef90 gene families (Suzek et al., 2015). For each input donor (fecal) metagenome, HUMAnN 3.6 provides bacterial-specific genes and pathway abundance whenever genes can be attributed to known bacterial species in the database. By default, gene families are annotated using UniRef90 definitions and pathways using MetaCyc definitions (Caspi et al., 2020). Whenever inferred genes cannot be attributed to bacterial species, gene and pathway abundances are reported without specifying their bacterial provenance. To identify metabolic pathways that are differentially abundant in patients or controls, we used linear discriminatory analysis with effect size estimation (LEfSe) implemented in the microbial R package. For pathways that showed strong enrichment in patients (pathways with the largest effect size, the LDA score), we also inferred bacterial species, genera, and higher-level taxa that were major contributors to that pathway.

## Results

### Baseline characteristics

A total of 23 untreated TB patients and 47 healthy controls were recruited in this study (Table S1). There was no significant difference in sex or age between the participating TB patients and HCs (p < 0.05, Table S1). The median age of the analyzed patients was 43 (IQR 34.5–55), and male TB patients significantly outnumbered females (61% vs. 39%). The patients were diagnosed with infiltrative tuberculosis (65.2%), disseminated tuberculosis (30.4%), and caseous pneumonia (4.3%).

### The gut microbiome composition in TB patients is strongly divergent from that in healthy controls

To obtain a comprehensive description of the gut microbiome, we searched for known (isolated) and unknown microbial taxons inferred from metagenome-assembled genomes (MAGs). Unlike traditional approaches that provide a truncated picture of the microbial diversity, this novel approach, implemented in Metaphlan 4.0 (Blanco-Míguez et al., 2023), provides a more complete description of the microbial content of the gut. We detected 987 microbial taxa in our combined set of treatment-naive TB patients and HCs. Our data show that gut community diversity (alpha-diversity) is not reduced in treatment-naive TB patients (Fig. 1A) but shows statistically significant divergence (Bray-Curtis dissimilarity) from the gut community in HCs (p = 0.00747) (Fig. 1B). We next showed that the extent of community divergence is strong enough to separate patients from controls in an unsupervised manner using principal component analysis (PCA). The principal component analysis clearly separated TB patients from HCs along the first PC, the strongest axis of microbial variation (Fig. 1C). Our results suggest that untreated TB patients arrive at the hospital with a strongly disturbed gut community.

**Figure 1.**
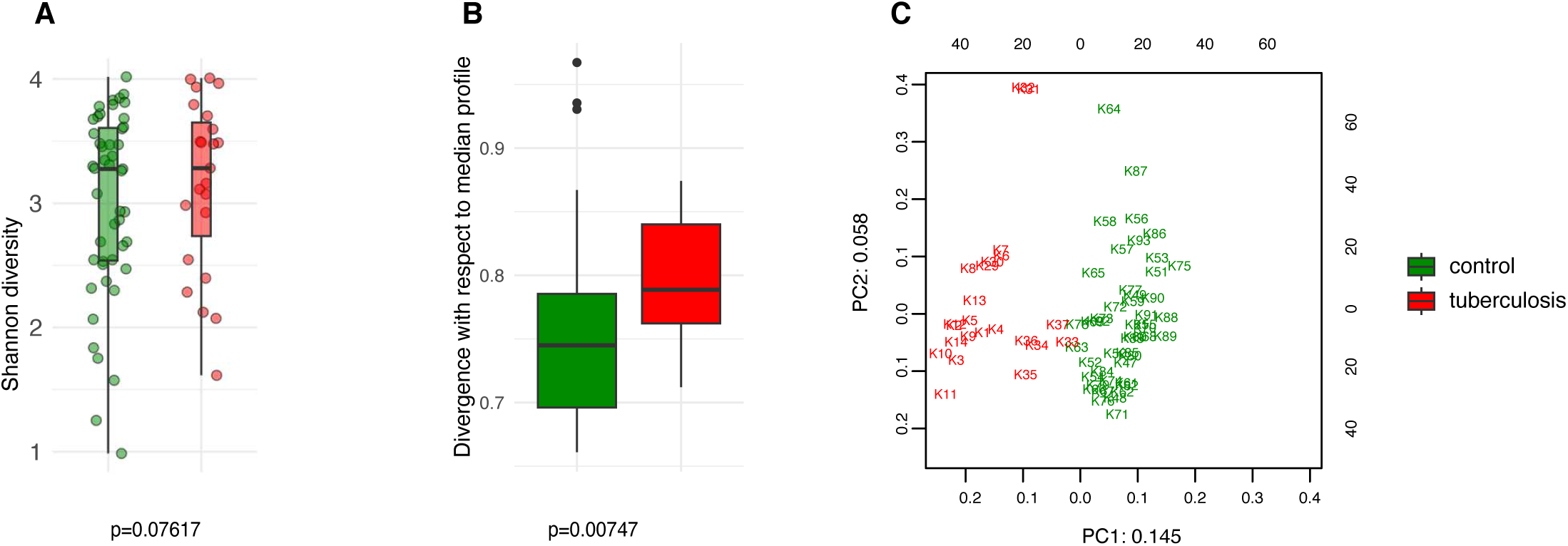
Comparison of the intestinal microbiota diversity in untreated TB patients versus healthy controls. **A** The alpha diversity was assessed using the Shannon index. No significant differences in diversity between the TB and control groups (P > 0.05); **B** The beta diversity, assessed as Bray-Curtis dissimilarity, indicates significant divergence between the TB patients and controls (P < 0.007); **C** PCA analysis based on CLR-transformed relative abundance. PC1 and PC2 are given with the proportion of total variance explained

### Differentially abundant bacterial taxa between TB patients and healthy controls

To facilitate comparison with published data, we characterized microbiomes at different taxonomic resolutions. At the phylum level, TB patients showed an apparent increase in *Firmicutes* abundance (36.4% vs. 21.3% in controls, p = 0.03) and an elevated proportion of *Actinobacteria* (1.8% vs. 0.2%, p = 0.0002) and *Proteobacteria* (5.6% vs. 3.8%, p > 0.05) (Fig. 2A). This increase in *Firmicutes* was at the expense of *Bacteroidetes* (54% vs. 72.2%), which is a frequent sign of a disturbed gut microbiome. Taxonomic composition at the genus, family, and order levels was also strongly altered in TB patients (Figure S1).

**Figure 2.**
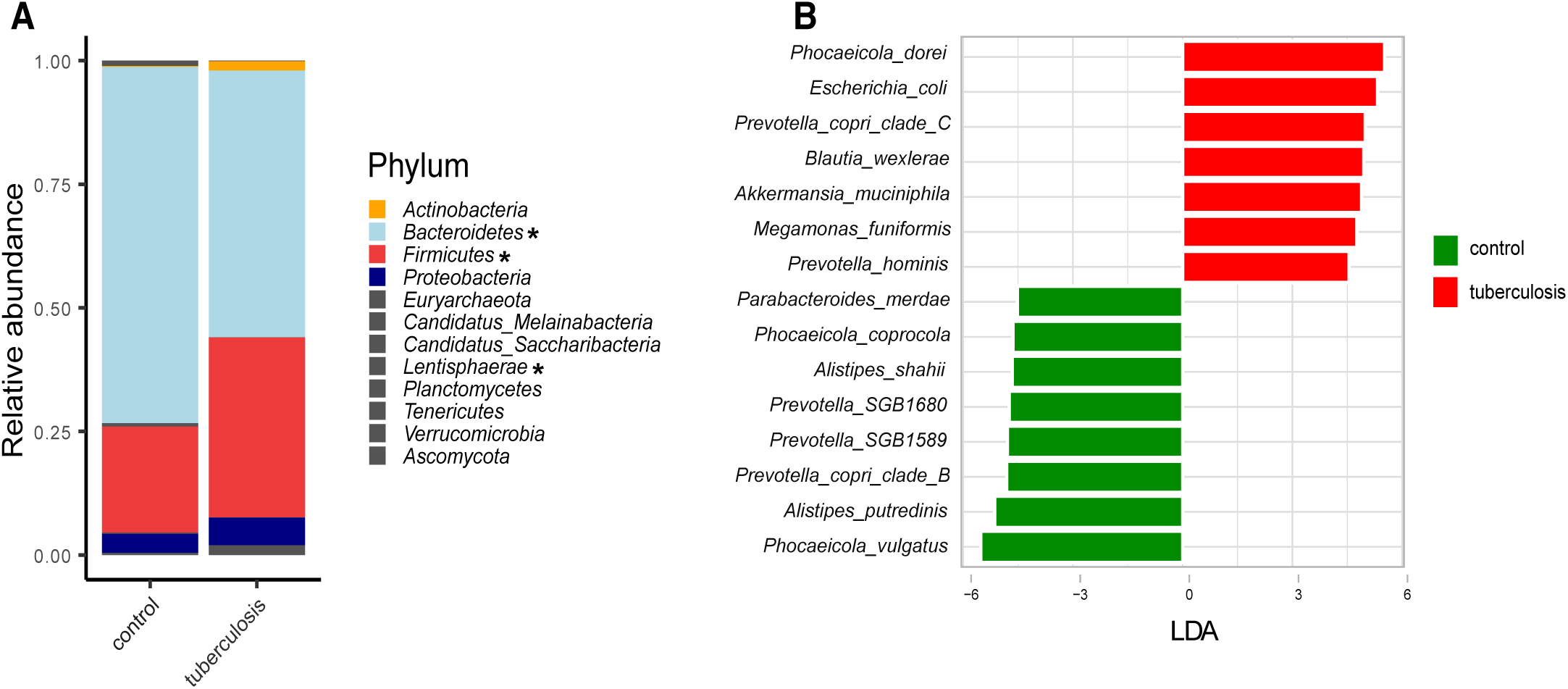
Taxonomic composition of the fecal microbiota of TB patients and healthy controls. **A** The average relative abundance of bacterial taxa at the phylum level in TB patients and HCs. Asterisks indicate taxa that showed statistically significant differences between the TB group and HCs based on the ANCOM statistical analysis (P < 0.05); **B** Barplot depicting top bacterial species (LDA scores > 4.5) discriminating between groups.

We next used LEfSe, a linear discriminant analysis, to identify microbial species that differentiate TB patients’ gut microbiomes from HCs (Segata et al., 2011). For each taxon, LEfSe also estimates the effect size, the LDA-score. Altogether, 205 taxa (species and subclades defined by MAGs) showed differential abundance (adjusted p-value ≤ 0.05 and LDA-score ≥ 2.5) (Supplementary Table 2), and we highlighted bacterial species with the strongest differentiation (LDA ≥ 4.5) (Fig. 2B). For example, *Phocaeicola dorei, Escherichia coli, Prevotella copri clade C,* and *Akkermansia muciniphila* were strongly enriched in TB patients (Fig. 2B). In contrast, *Phocaeicola vulgatus, Alistipes putredinis, Prevotella copri clade B,* and *Prevotella SGB1589* were more abundant in the HCs (Fig. 2B). The top TB-associated taxon, *Phocaeicola (Bacteroides) dorei,* was previously isolated from patients with bacteremia (Cobo et al., 2022). Despite wide interest in the role of *P. (B.) dorei* in the intestinal microbiome, this bacterium is poorly studied, and there is little evidence of its direct involvement in the pathogenesis of infections (Cobo et al., 2022; Yoshida et al., 2018). Its accurate identification is challenging due to its close relationship with *P. vulgatus* (96% similarity), one of the most numerically predominant *Bacteroides* species in the human intestine (Pedersen et al., 2013; Vu et al., 2022). Unlike *P. (B.) dorei*, other top TB-associated species, such as *E. coli, A. muciniphila*, and *M. funiformis,* are relatively better studied and were previously implicated in various diseases, such as inflammatory bowel diseases (Crohn’s disease and colitis), metabolic syndromes (obesity, hypertension, type 2 diabetes), autism, and infections (diarrhea, cystitis) (Sheng et al., 2022; Zou et al., 2020).

We next inspected the abundance of these top discriminatory species. We noted that some healthy donors carried high proportions of TB-associated species, such as *Phocaeicola dorei, Escherichia coli*, and *Prevotella copri clade C* (Fig. 3A, asterisk). Our bacterial culturing test for these healthy donors showed overgrowth of *C. albicans* or lactose-negative *E. coli,* and the coprogram stool test showed iodophilic flora, a sign of an imbalance between normal and pathogenic gut flora (our unpublished data). Hence, these healthy outliers and published data suggest that the top TB-associated species are pathobionts that correlate with intestinal dysbiosis and poor health. Notably, these potential pathobionts comprise a quarter of the gut microbiome in some TB patients (shown in orange in Figure 3B), a high toll on the host immunity given the strong deficit in health-associated commensals.

**Figure 3.**
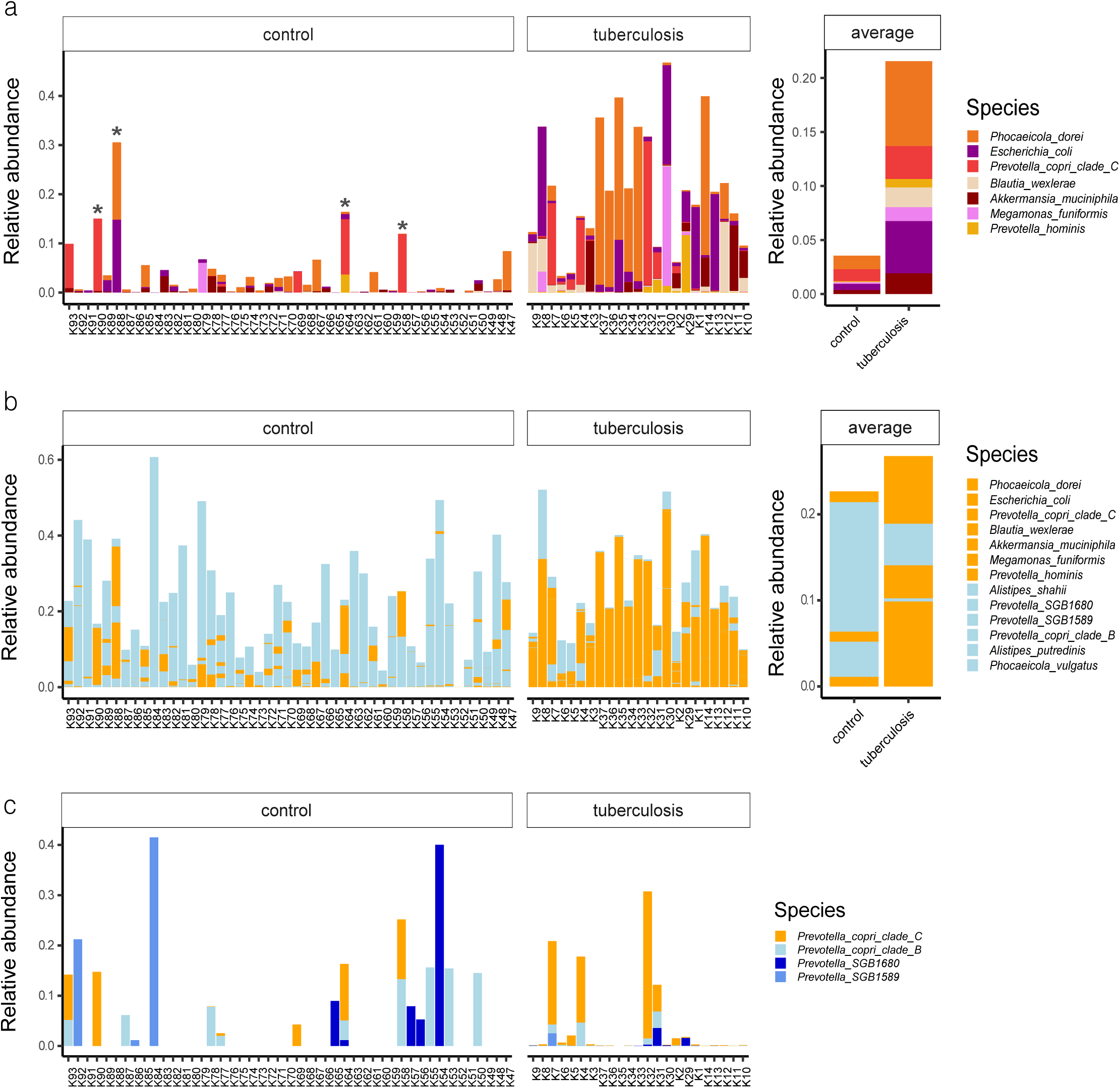
The relative abundances of top TB-associated species in the gut microbiota of TB patients and healthy controls. **A** The relative abundances of top TB-associated species (LDA scores > 4.5) in TB patients and HCs lined up on the x-axis with respective IDs. Asterisks indicate outlier healthy donors carrying a high proportion of TB-associated species; Averaged abundances are shown on the right-hand side barplot; **B** Combined relative abundances of TB-associated species (potential pathobionts) and health-associated commensals in donors. In orange potential pathobionts, in light-blue – health-associated commensals; **C** The distribution of *Prevotella copri* clades and related lineages in the gut microbiota of TB patients and healthy controls. *Prevotella copri clade C* in orange was associated with TB, while clades in blue were more prevalent in healthy donors.

### Depleted strain diversity of common gut commensals in TB patients

We used strain-level profiling with MetaPhlAn 4 (Blanco-Míguez et al., 2023) and uncovered subspecies diversity that showed contrasting enrichments in TB patients and HCs. For example, according to linear discriminant analysis, *Prevotella copri clade C* was enriched in TB patients, while *Prevotella copri clade B* was associated with controls (Fig. 2B, Supplementary Table 2). Closer inspection revealed that healthy donors featured a more diverse collection of *Prevotella* subclades and lineages than TB patients (Fig. 3C). We looked for published data to see if *Prevotella copri* clades are associated with human diseases. While earlier studies reported conflicting correlations between *Prevotella copri* and various diseases (Dillon et al., 2014; H. K. Pedersen et al., 2016), a recent large-scale study did not find evidence for these earlier claims (Tett et al., 2019). Instead, Tett and colleagues found that *Prevotella copri* clade diversity was reduced in much of the world’s populations with Westernized diets and disease settings, while non-industrialized societies with fiber-rich traditional diets featured richer diversity (Tett et al., 2019). Our study shows that TB patients show reduced diversity of *Prevotella copri* clades. We, therefore, hypothesize that *Prevotella copri clade C’s* association with TB could be driven by a depletion of other lineages (Fig. 3C) rather than the detrimental role of this particular clade.

### The metabolic potential of the gut microbiota in TB patients points to increased fermentation and lactic acidosis

The microbial community constantly adapts to the intestinal environment by increasing or decreasing certain species and, hence, certain gene assemblages and metabolic pathways. We inferred metabolic pathways via gene assemblages in the analyzed microbiomes using the HUMAnN 3.6 tool (Beghini et al., 2021). This approach allowed us to delineate quantitative changes in patients’ metabolic potential of the analyzed gut microbes. In total, 131 metabolic pathways were differentially enriched between TB patients and controls’ gut microbiomes (adjusted p-value ≤ 0.05 and LDA-score ≥ 2). Notably, 120 pathways were overrepresented in TB patients’ microbiomes (p ≤ 0.05, Supplementary Table 3, Fig. S3), as compared to only 11 in HCs. When clustered into higher-level MetaCyc pathway groups, this profound increase in metabolic capacity revealed an elevation in biosynthetic processes supporting microbial cell division and growth (Figure 4A, C). This net increase in pathways supporting bacterial growth in patients’ gut contrasts with the metabolic profile in healthy donors, where metabolic pathways reflect the normal functioning of the bacterial cell (Fig. 4A).

**Figure 4.**
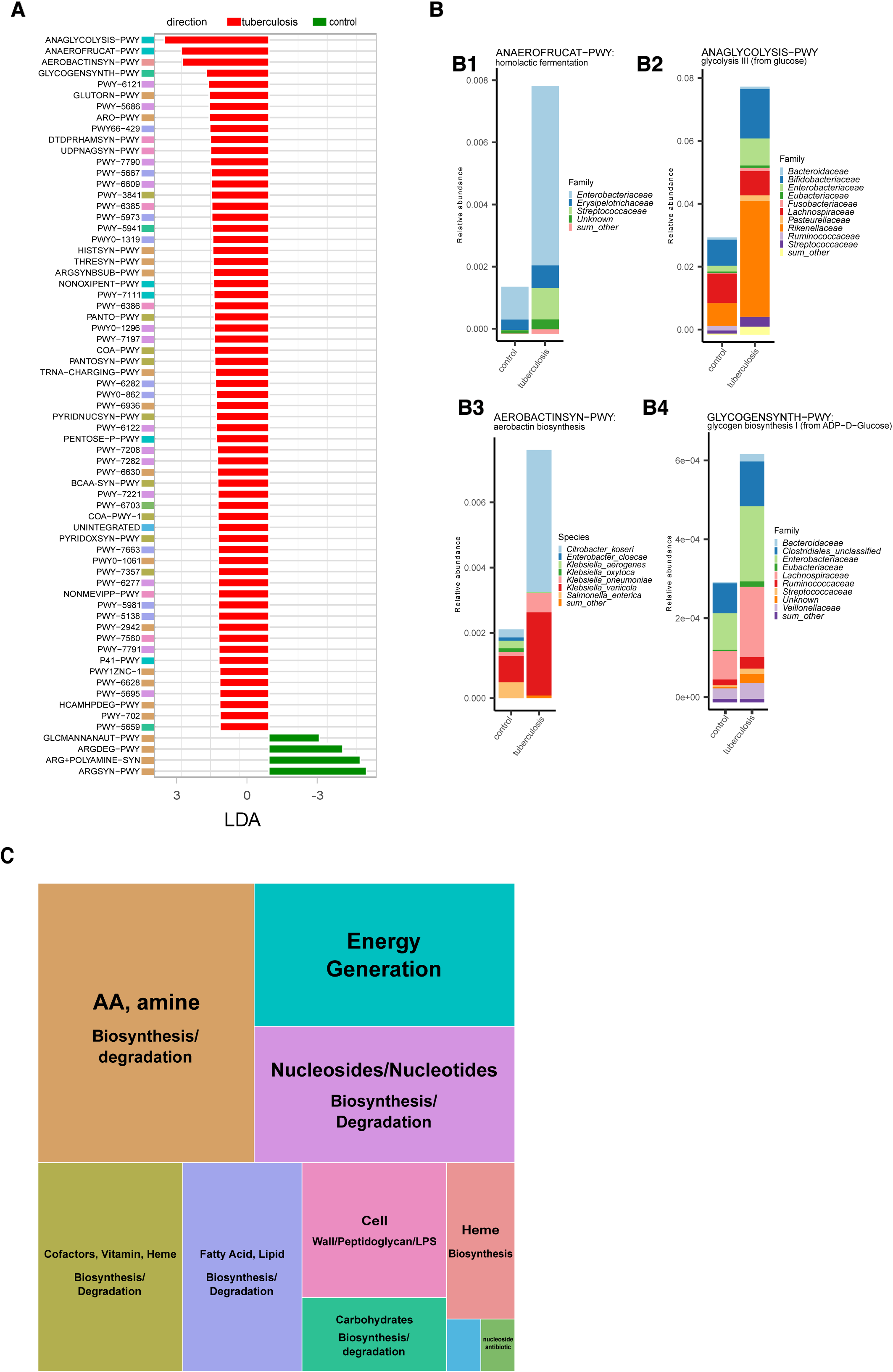
Microbial metabolic pathways increased in TB patients. **A** Metabolic pathways were differentially enriched (LDA score > 2,5) in TB patients and healthy controls based on linear discriminant analysis. Metabolic pathway names represent MetaCyc identifiers. Full pathway names are given in Fig. S3. Colored boxes for each metabolic pathway indicate membership in the higher-level MetaCyc pathway types summarized in Fig. 3C; **B** Abundance of the four metabolic pathways enriched in TB patients and contribution of bacterial taxa; The total height of the stacked barplot corresponds to the fraction of the pathway from the total amount of pathways in the gut microbiome in TB patients. Each colored section of the stacked barplot represents different bacterial families’ contributions to this pathway. Thus, colored sections in B1 show the contribution of bacterial families to homolactic fermentation, in B2 to glycolysis III (from glucose), in B4 to glycogen biosynthesis I (from ADP−D−Glucose); in B3, sections represent bacterial species contributions to the aerobactin biosynthesis pathway; **C** Summary of metabolic pathway types enriched in TB patients. The block size reflects the number of unique metabolic pathways belonging to a particular type of metabolic pathway.

Microbial metabolism and growth in the anaerobic colon can be achieved by different strategies for obtaining energy and biosynthetic precursors. The choice of the strategy depends on the carbon source, its oxidation state, pH, and other factors in the intestine (Wolfe Alan J., 2005). Hence, dominant strategies can be informative about the changes in the colon environment linked to the pathology. We, therefore, highlighted major changes in energy production pathways observed in the TB patients’ gut microbiome. Our analysis showed that the gut microbiome in TB patients is characterized by an increased presence of metabolic pathways aimed at glucose consumption and fermentation with end products such as acetate and lactate. Specifically, glycolysis (ANAGLYCOLYSIS-PWY), homolactic fermentation (ANAEROFRUCAT-PWY), pentose phosphate pathway (NONOXIPENT-PWY and PENTOSE-P-PWY), and pyruvate fermentation to acetate and (S)-lactate I (P41-PWY) were the top changes in TB patients gut microbiome (Fig. 4 A, B1, and B2). Higher use of glucose oxidation (glycolysis) compared to the HC microbiome can indicate higher availability of glucose in patients’ intestines. Glycolysis supports a higher rate of generating ATP and biosynthetic precursors for dividing bacteria. Relatively higher presence of the pentose phosphate pathway also points to higher glucose consumption, and reactions in this pathway synthesize additional biosynthetic precursors needed for dividing cells (Folch et al., 2021). The source of the higher glucose in the patient’s large intestine is unclear. Under homeostatic conditions, a small amount of simple sugars may reach the colon from the small intestine. Locally, simple sugars arise from host– and diet-derived complex carbohydrates fermented mostly by *Bacteroidetes* and are readily consumed by resident bacteria. In patients, however, we find depletion of *Bacteroidetes* and no increase in pathways degrading complex carbohydrates to explain higher amounts of simple sugars (Fig. S3, Supplementary Table 3).

A higher flux of glucose and increase in glycolysis in anaerobic conditions and pH in the normal range (5.5-7.5 in the large intestine lumen and 7.1-7.5 on the mucosal surface) (Nugent et al., 2001) usually leads to excretion of acetate (acetogenesis), ethanol, and format. However, our analysis suggests a relatively high fermentation to lactate and acetate (P41-PWY, PWY-5100, P461-PWY, and ANAEROFRUCAT-PWY) compared to HCs gut microbiome (Fig. 4A, 4 B1, Supplementary Table 3). While acetate and lactate are consumed by host cells and cross-feeding bacteria, acetate is normally the most abundant among short-chain fatty acids (SCFA) in the colon (Wolfe Alan J., 2005). In contrast, lactate is normally kept at very low concentrations by both host cells and lactic acid-degrading bacteria and has a higher potential to acidify the intestine (Louis et al., 2022). It is, therefore, notable that in TB patients’ gut microbiomes, lactic acid fermentation (homolactic fermentation) is among the top elevated pathways and may indicate lactic acidosis (Fig. 4 B1).

In line with our inference of higher bacterial growth in TB patients’ intestines, we observed increased biosynthetic pathways supporting cell wall synthesis (Fig. S4, Supplementary Table 3). For example, we identified an increase in pathways producing cell wall components such as peptidoglycan, lipopolysaccharide, and the enterobacterial common antigen (DTDPRHAMSYN-PWY; UDPNAGSYN-PWY; PWY-6386; PWY-7315; PWY0-1241; PWY-1269; PWY0-1586) (Fig. S4). For these pathways, we computed each bacterial genus and family’s contribution to the total estimate. This information allowed us to learn about bacterial taxa assembling cell walls at an increased pace (Fig. S4). Our analysis identified major contributors (taxa contributing more than 1% of the total) from the following families: *Enterobacteriaceae, Streptococcaceae, Desulfovibrionaceae, Lachnospiraceae, Ruminococcaceae, Eubacteriaceae, Clostridiales, Prevotellaceae, Selenomonadaceae, Hafniaceae, Bacteroidaceae,* and *Vibrionaceae,* belonging mainly to the phylum *Proteobacteria, Firmicutes,* and *Bacteroidetes*. Notably, several bacterial families, such as *Enterobacteriaceae, Streptococcaceae, Eubacteriaceae,* and others, that contributed to higher cell wall synthesis pathways also contributed to the higher presence of pathways for energy production through glycolysis and homolactic fermentation (Fig. 4 B1-B4).

## Discussion

Previous studies characterized taxonomic changes in TB patients’ gut microbiomes; however, how these changes influenced host immunity was unclear and required insight into microbial function. We used deep sequencing and showed that gut community alterations in TB patients are accompanied by increased metabolic pathways leading to cell division and growth. To interpret these metabolic shifts further, we paid special attention to pathways of energy acquisition, which, on the one hand, informs on the colon environment and, on the other hand, on the metabolic output of the community and, hence, on their potential impact on the host. We found that the patient’s gut microbes were characterized by increased glucose consumption via glycolysis and lactate fermentation, which hinted at an unhealthy colon environment compared to healthy donors (Fig. 3A, C).

Higher use of glycolysis presumably indicates higher than usual glucose availability in the TB patient’s colon, which is a potent cue for most microbes to switch the metabolic program (Wolfe, 2015). The source of higher glucose in patients’ large intestines is unclear. Normally, in the large intestine, the majority of available carbohydrates that the host does not consume are diet– (i.e., fiber from plant foods) or host-derived (e.g., mucin, cellular debris) complex carbohydrates (polysaccharides); they are broken down by dominant residents of the colon, primarily *Bacteroides* and *Clostridiales* (Kamada et al., 2013). However, our taxonomic analysis showed that *Bacteroides* were depleted in TB patients. In contrast, *Proteobacteria (Enterobacteriaceae)*, such as *E. coli*, which cannot normally break down polysaccharides into simple carbohydrates, increased (Fig. 2B and Fig. 4 B1). Not surprisingly, *Clostridiales*, which can consume complex and simple carbohydrates, were increased (Fig. S2).

Irrespective of the source, a higher flux of glucose causes most bacteria to prefer glycolysis to grow and divide rapidly (Wolfe, 2015). Indeed, glycolysis supports 110 times faster generation of biosynthetic precursors and energy in terms of ATP but rapidly produces lactate (Wolfe, 2015). Using species-specific marker genes, we identified that increased glycolysis (glycolysis III from glucose) in patients’ guts was explained by taxa that generally contain harmful species. Namely, *Enterobacteriaceae* (*E. coli, Citrobacter freundii*, and *Klebsiella pneumoniae*), *Streptococcaceae (Streptococcus thermophilus, S. salivarius, S. parasanguinis, S. gallolyticus), Rikenellaceae* (*Alistipes dispar, A. communis*), and *Bifidobacteriaceae* (*Bifidobacterium longum, B. catenulatum, B. pseudocatenulatum*, and *B. breve*) (Fig. 4 B2).

Our findings show that homolactic acid fermentation is the next top shift in the gut microbiome metabolism in patients, which is consistent with the higher glycolysis. Taxa featured in higher glycolysis were among the major contributors to increased capacity for homolactic fermentation. Namely, representatives of *Enterobacteriaceae* (e.g., *E. coli, C. freundii,* and *K. pneumoniae*) and *Streptococcaceae (e.g., Streptococcus thermophilus, S. salivarius, S. parasanguinis,* and *S. gallolyticus*), and *Erysipelotrichaceae (Clostridium innocuum*) (Fig. 4 B1). Elevated production of lactate and acetate is evidenced by an increase in other pathways producing these acids (Fig. 4A). Our findings, namely increased capacity for lactic acid fermentation and major contributing taxa, are consistent with the experimental data showing that homolactic acid fermentation is used by lactic acid bacteria, such as *Lactobacillus, Lactococcus*, and many *Streptococci,* as well as members of the family *Enterobacteriaceae*. *Enterobacteriaceae*, in addition, can perform mixed-acid (heterolactic) fermentation.

The higher presence of pathways for lactic acid fermentation and glycolysis indicates a higher propensity of the gut community to acidify the intestine (Wolfe, 2015; Kamada et al., 2013). While the host tissue can absorb lactate, the gut community also responds by increasing the number of lactate-utilizing bacteria (Russell & Rychlik, 2001; Louis et al., 2022). We examined our taxonomic data to test this predicted compensatory increase in lactate-utilizing species. Our taxonomic data shows that several prominent lactate utilizers are strongly increased in TB patients at species or genus levels. Specifically, *Coprococcus catus* (Louis et al., 2022) and *Megasphaera BL7* (Shetty et al., 2013), which use the acrylate pathway, were strongly increased in TB patients (Suppl. Table 2). At the genus level, TB patients show enrichment of the two highly adapted lactate-utilizing genera, *Anaerobutyricum* and *Anaerostipes* (Fig. S2). While these observations are consistent with a compensatory increase in lactate utilizers, counting the abundance of lactate utilization pathways is not straightforward. Thus, the acrylate pathway is used only by a limited number of human gut bacteria, and most other lactate-utilization pathways involve conversion to pyruvate. Pyruvate makes lactate utilization intertwined with central metabolic pathways that can start with any major organic compound, i.e., carbohydrates, lipids, and amino acids. Nevertheless, the only testable acrylate pathway did not feature in patients according to linear discriminant analysis (Figure 4A) (Reichardt et al., 2014). We recognize that a more advanced approach might disentangle this complexity. However, our current methodology renders the accurate assessment of lactate utilization challenging, which is a limitation of current work.

The accumulation of lactate in the colon decreases pH and strongly inhibits the resident commensal flora, especially *Bacteroidetes* (S. P. Wang et al., 2020). Our taxonomic analysis showed a systematic decrease in *Bacteroidetes* proportion in patients and an increase in *Actinobacteria, Proteobacteria,* and *Firmicutes* (Figure 2A). Observed taxonomic changes closely mirrored the effect of lactate acidosis, as demonstrated by experiments and mathematical modeling. Thus, experiments that modeled the effect of pH and lactate on human colon microbiota demonstrated that lactate acidosis strongly depletes *Bacteroidetes* and increases *Actinobacteria, Proteobacteria,* and some *Firmicutes*, such as lactobacilli, well-known lactate producers that are tolerant to low pH (Wang et al., 2020). This experiment showed that lactic acidosis destabilized resident flora and promoted pathogenic bacteria, such as *Streptococcus bovis, Salmonella, E. coli, Campylobacter jejuni, C. difficile,* and *Vibrio cholerae,* that are adapted to low pH and get advantage of lactate (Louis et al., 2022). Our taxonomic analysis also showed an increased abundance of similar or closely related pathobionts in TB patients, such as *E. coli, C. freundii*, *K. pneumoniae*, *C. innocuum,* and different species in *Streptococcaceae,* that are associated with diverse human diseases (Chhatwal & Graham, 2017; Crum-Cianflone, 2009; Li et al., 2022; Santos et al., 2020; Xue et al., 2023). Moreover, the combined relative abundance of these pathobionts in patients reached 1/4 of the total microbiome content (Figures 3A and 3B). The extent of the discovered pathobiont burden likely poses a heavy toll on host immunity on top of the mycobacterium infection.

Since our metabolic reconstructions suggest lactate acidosis, we conclude by discussing lactate’s direct effects on the course of MTB infection and TB treatment outcome. In clinical tuberculosis, the balance between the antibacterial activity of macrophages and tissue destruction by extracellular enzymes, primarily matrix metalloproteinases (MMPs), produced during inflammation is critical to disease outcome and transmission (Ong et al., 2014). Recent studies show that this balance is affected by the extracellular concentration of lactate (Whittington et al., 2023), which is produced by infected macrophages using glycolysis (Langston et al., 2017). Mechanistically, lactate, on the one hand, improves clearance of MTB in the already infected macrophage by promoting autophagy. On the other hand, it markedly suppresses both TNF-α and IFN-γ but leaves IL6 and IL10 unaffected, thereby interfering with further enhancement of anti-MTB activity (Ó Maoldomhnaigh et al., 2021). Moreover, lactate specifically upregulates matrix metalloproteinases that cause lung destruction in tuberculosis (Whittington et al., 2023), the process known to result in higher morbidity and mortality in TB treatment (Ong et al., 2014). Thus, recent mechanistic insights underscore the multifaceted effect of lactate on Th1 response with an overall negative impact on the ability to cope with MTB, which is more in line with clinical data. Indeed, serum lactic acidosis is associated with higher mortality during TB treatment (Im et al., 2021; Ntambwe & Maryet, 2012) and other infections (Trzeciak et al., 2007), and monitoring lactate acidosis was shown to help rescue TB patients (Im et al., 2021).

In the context of ongoing inflammation, high lactate in patients serum can also be attributed to the glycolytic activity of immune cells and an inflamed tissue environment (Manosalva et al., 2021). Our findings point to an additional unacknowledged source for high lactate in the patients’ sera, the gut microbiota. Thus, in light of lactate’s importance in disease outcome, our findings point to an intriguing avenue to understand the role of gut microbiota in TB treatment efficacy. Namely, collecting clinical data along with serum and gut microbiome readouts, such as fecal metabolites and deeply sequenced metagenomes, to directly assess sources of intestinal lactate acidosis, its possible impact on serum levels, and TB outcome. Importantly, the idea to target lactic acidosis in TB treatment as a new host-directed therapy was proposed earlier by others (Kiran & Basaraba, 2021). In animal studies, excess fermentation in the gut has been long known to cause lactic acidosis and treatment measures are actively studied (Nagaraja & Titgemeyer, 2007). Here, we add that one major source of lactic acidosis can be intestinal microbiota-driven acidosis.

## Acknowledgements

This research was supported by the Russian Science Foundation (grant 22-25-00272). The foundation had no role in data collection, data analysis, data interpretation, or writing of the report.

## Author сontributions

MY, LB and BY conceived and designed the study. LB, AZ, FB, MY and SB were responsible for the selection of patients for the study, provided clinical services, extracted DNA from feces, and collected study data. LB and MY supervised the study. DT, AB, FB, RA, and BY analyzed the data. BY, FB and RA were responsible for the methodology and provided statistical expertise. MY and BY interpreted the results and drafted the manuscript. MY, LB, AB, DT, FB, and BY contributed to the writing of the manuscript. MY and BY took care of the editing of the manuscript and the final approval of the version for publication. All the authors have read and approved the final manuscript.

## Ethical approval

This study was approved by the Ethics Committee of the Republican Clinical Antituberculous Dispensary (Russia). All subjects provided informed consent for this study. This project was performed in accordance with the approved guidelines based on the ethical principles outlined in the Declaration of Helsinki.

## Data sharing

The metagenomic sequencing data will be deposited to the NCBI Sequence Read Archive (SRA) immediately following publication and ending 36 months following article publication. The data can be provided for individual participant data meta-analysis. Proposals should be directed to the corresponding author

## Additional information

Supplementary information

Supplementary Table 2.

Supplementary Table 3.

