## Supplementary Information for "Excess fermentation and lactic acidosis as detrimental functions of the gut microbes in treatment-naive TB patients"

Table of Contents

Table S1. Clinical and demographic characteristics of the compared groups	2

Figure S1. Taxonomic composition of the fecal microbiota of TB patients and healthy controls	3

Figure S2. Taxonomic features of the gut microbiota of TB patients and controls	4

Figure S3. Metabolic features of the gut microbiota of TB patients and controls	5

Figure S4. Metabolic pathways of cell wall biosynthesis of the gut microbiota of TB patients and controls	6

**Table S1. Clinical and demographic characteristics of the compared groups**

|  | **TB patients**  **(N =23)** | **Healthy control**  **(N =47)** |
| --- | --- | --- |
| Female | 9/23 (39%) | 22/47 (47%) |
| Men | 14/23 (61%) | 25/47 (53%) |
| Age, years (Median (IQR) | 44 (38-53) | 40 (36-45) |
| Clinical forms of pulmonary tuberculosis | |  |
| Infiltrative TB | 15/23 (65,2%) | .. |
| Disseminated TB | 7/23 (30,4%) | .. |
| Caseous pneumonia | 1/23 (4,3%) | .. |

Data are median (IQR) (first and third quartiles) or n/N (%), unless stated otherwise. N ― number of individuals

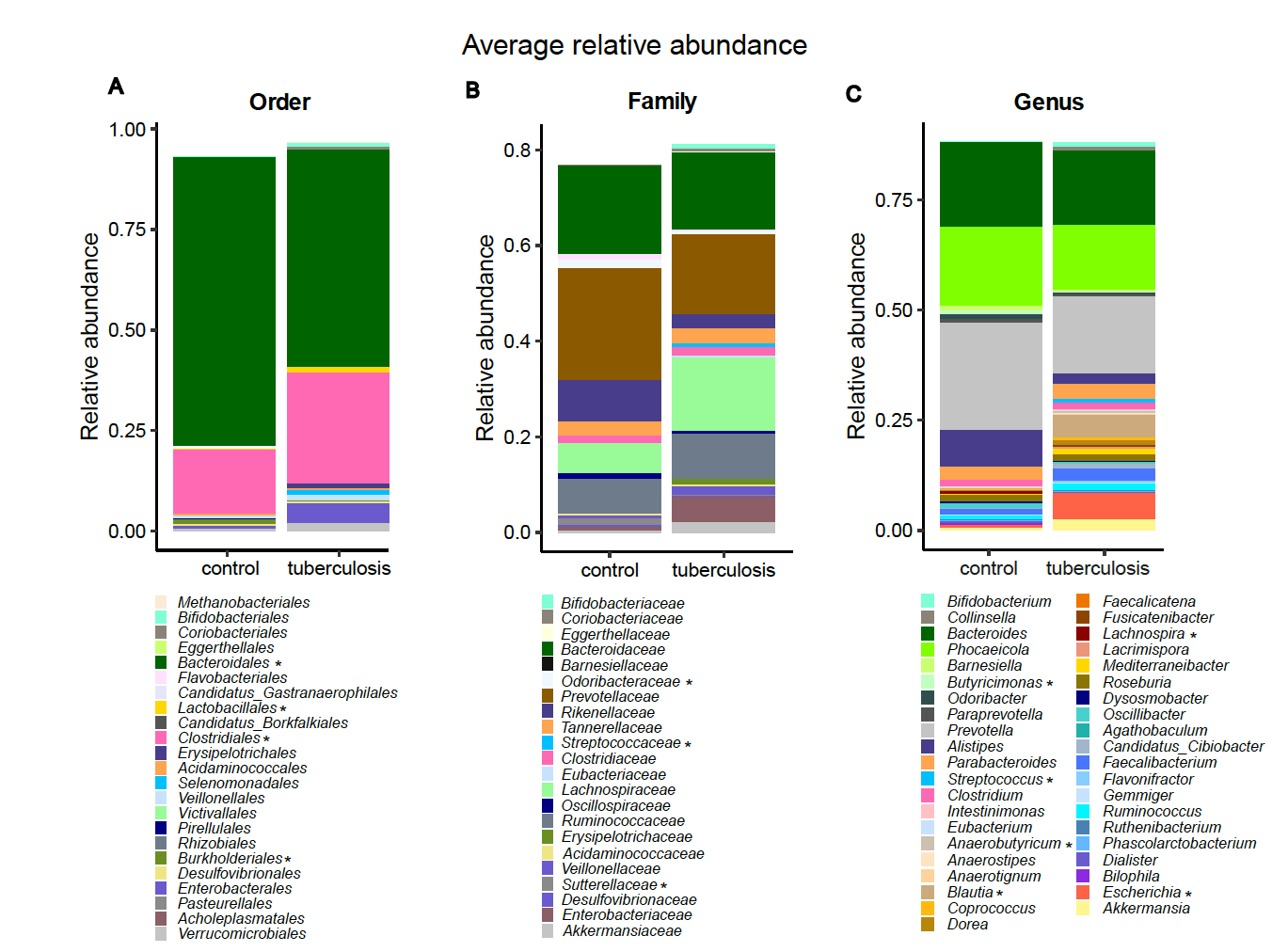

**Figure S1. Taxonomic composition of the fecal microbiota of TB patients and healthy controls.**

Plots show the average relative abundance of microbial taxa in the gut microbiota of TB patients and controls at the order (A), family (B), and genus (C) levels. A statistical difference in proportions between groups was tested using the ANCOM statistical framework. An asterisk indicates taxa that are statistically significantly different between the compared groups (P < 0,05). We required bacterial taxa to be present in 60% of donors at an abundance of 5% or more. In addition, we removed unclassified taxa (such as FGB, GGB, etc.) for the visualization. Taxonomic composition at the genus, family, and order levels is strongly altered in TB patients.

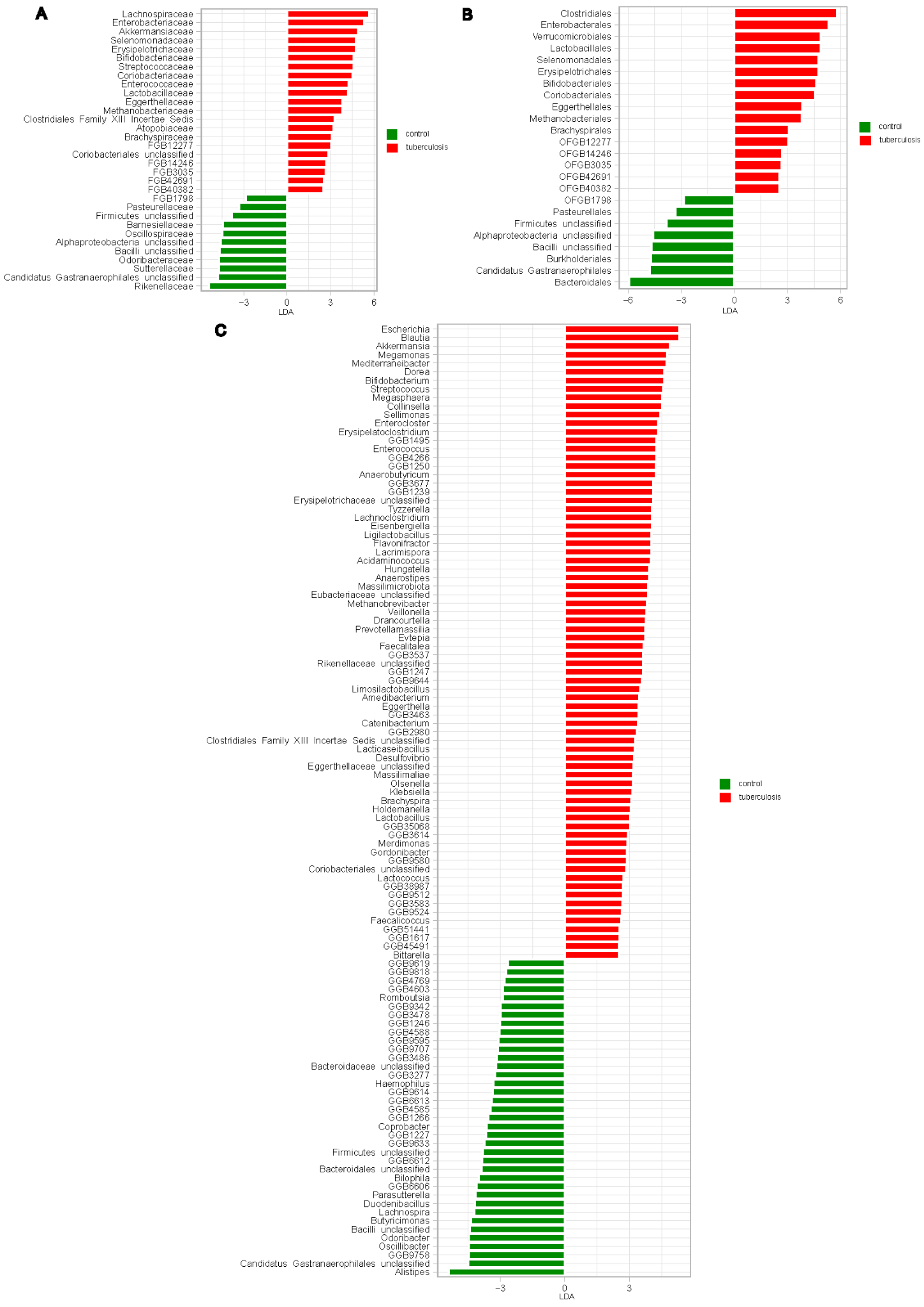

**Figure S2. Taxonomic features of the gut microbiota of TB patients and controls.**

LDA score plots were generated using the LEfSe analysis. The length of the bar column represents the LDA score. Plots show the microbial taxa with significant differences between the TB patients (red) and healthy controls (green) at the order (A), family (B), and genus (C) level (LDA score > 2,5). Two prominent lactate utilizers that use the acrylate pathway, *Coprococcus catus* and *Megasphaera BL,* were strongly increased in TB patients. Family *Clostridiales*, which can consume both complex and simple carbohydrates, were increased in TB patients.

.

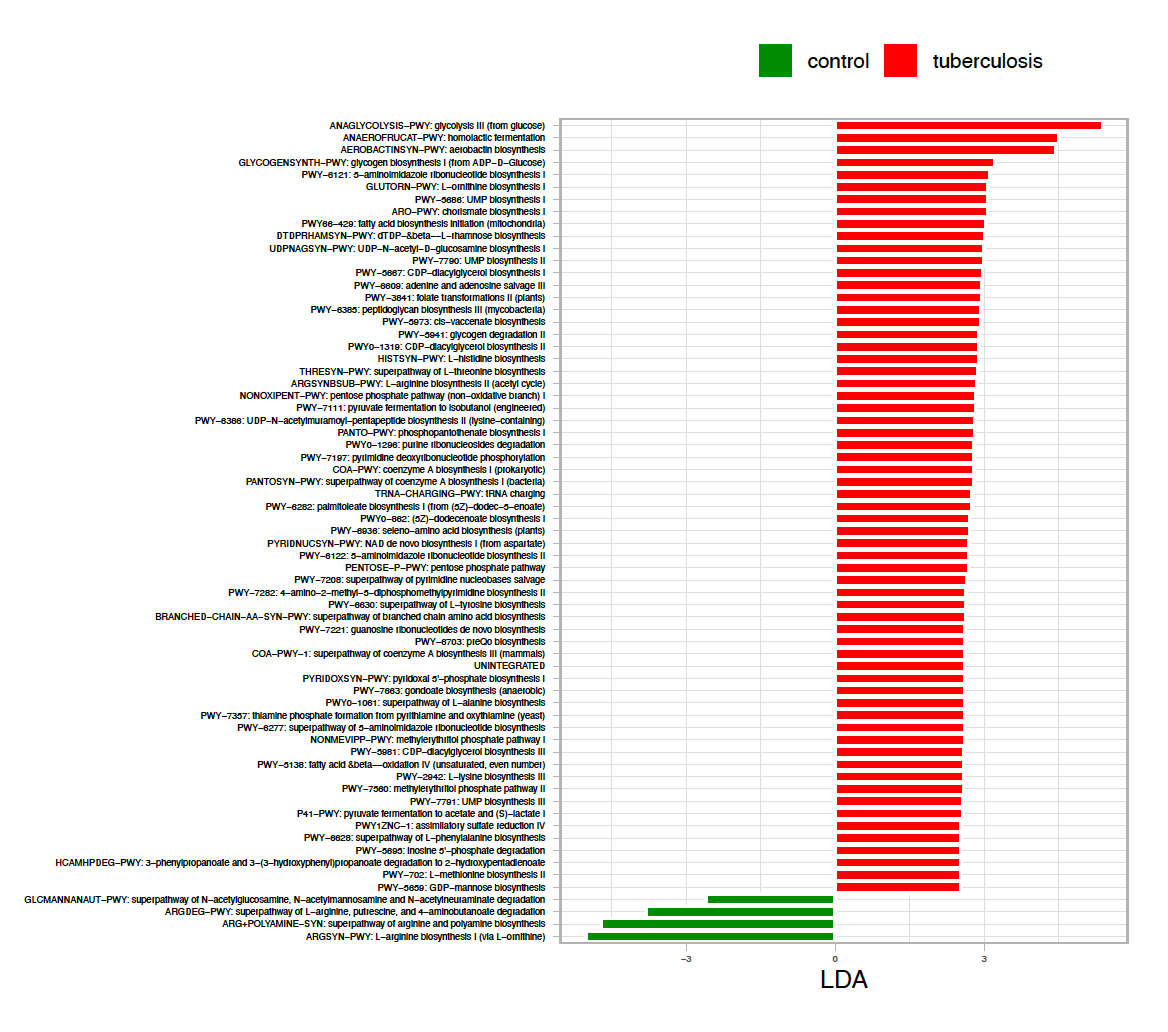

**Figure S3. Metabolic features of the gut microbiota of TB patients and controls.**

LDA score plots were generated using the LEfSe analysis. The length of the bar column represents the LDA score. Plots show metabolic pathways with significant differences between the TB patients (red) and healthy controls (green) (LDA score > 2,5). Metabolic pathways are inferred based on HUMAnN 3.6 and annotated based on Metacyc. In the gut microbiota of TB patients, there is an increase in the number of pathways supporting bacterial growth, while the metabolic profile of healthy donors reflects normal bacterial cell function.

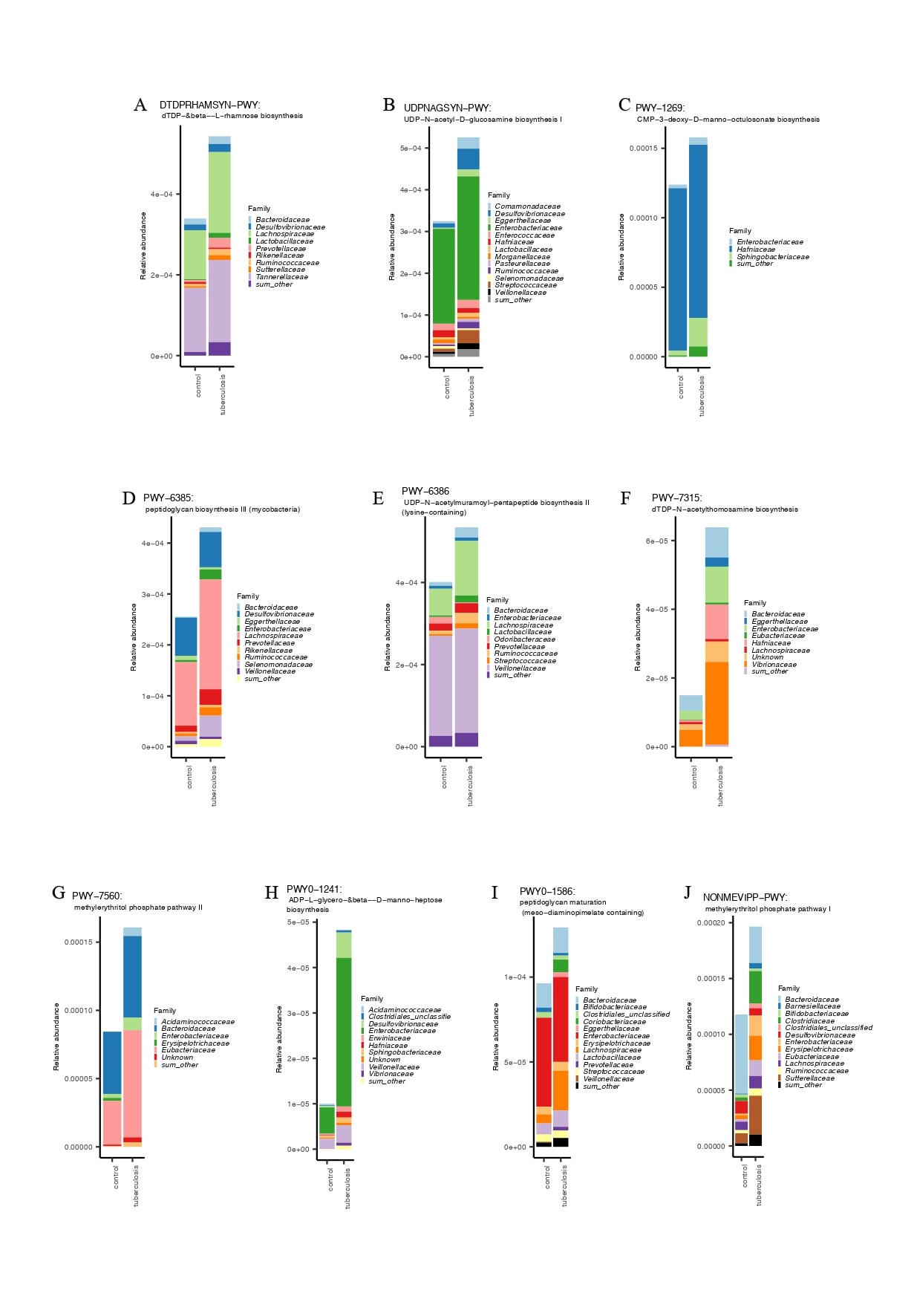

**Figure S4. Metabolic pathways of cell wall biosynthesis of the gut microbiota of TB patients and controls.**

A - Relative abundance of dTDP-&beta--L-rhamnose biosynthesis pathway in the gut microbiota of TB patients and controls; B - Relative abundance of UDP-N-acetylmuramoyl-pentapeptide biosynthesis II (lysine-containing) pathway in the gut microbiota of TB patients and controls; C - Relative abundance of CMP-3-deoxy-D-manno-octulosonate biosynthesis pathway in the gut microbiota of TB patients and controls; D - Relative abundance of peptidoglycan biosynthesis III (mycobacteria) pathway in the gut microbiota of TB patients and controls; E - Relative abundance of UDP-N-acetylmuramoyl-pentapeptide biosynthesis II (lysine-containing) pathway in the gut microbiota of TB patients and controls; F - Relative abundance of dTDP-N-acetylthomosamine biosynthesis pathway in the gut microbiota of TB patients and controls; G - Relative abundance of methylerythritol phosphate pathway II in the gut microbiota of TB patients and controls; H - Relative abundance of ADP-L-glycero-&beta--D-manno-heptose biosynthesis pathway in the gut microbiota of TB patients and controls; I -Relative abundance of peptidoglycan maturation (meso-diaminopimelate containing) pathway in the gut microbiota of TB patients and controls; J - Relative abundance of methylerythritol phosphate pathway I in the gut microbiota of TB patients and controls. The plots show the relative abundance of metabolic pathways in TB patients and healthy donors and the contribution of bacterial taxa. Metabolic pathways are inferred based on HUMAnN 3.6 and annotated based on MetaCyc. The TB patient group has a higher abundance of metabolic pathways related to cell wall synthesis than the control group.
